# “Regressing Away Common Neural Choice Signals does not make them Artifacts. Comment on Frömer et al (2024, Nature Human Behaviour)”

**DOI:** 10.1101/2024.09.26.614447

**Authors:** Redmond G. O’Connell, Elaine A. Corbett, Elisabeth Parés-Pujolràs, Daniel Feuerriegel, Simon P. Kelly

## Abstract

Frömer et al (2024, Nature Human Behaviour) apply a deconvolution method to correct for component overlap in the event-related potential. They report that this method eliminates signatures of sensory evidence accumulation from response-aligned measurements of the centro-parietal positivity (CPP), suggesting that these signatures arise artifactually.. Here, we argue that the analysis and interpretation of their perceptual choice data are critically flawed. We demonstrate with simulations that the deconvolution analyses used by the authors are not designed to reliably test for the presence or absence of bounded accumulation signals.

## Main Text

Frömer et al ^1^ describe an experiment devised to identify neural correlates of value-based choice in the human event-related potential (ERP). Their analyses focus on a component known as the centro-parietal positivity (CPP) that has been widely implicated in tracing the sensory evidence accumulation (EA) process underpinning perceptual decisions ^2^ but has rarely been examined in the context of value-based decisions. Frömer et al’s paper makes a valuable contribution in providing compelling evidence, from response-locked waveforms plotted by response time (RT), that the CPP does not trace the value-based choice process that culminates in action selection.

In addition to their analysis of the value-based choice data, the authors analyzed ERPs from three pre-existing perceptual choice datasets. In these datasets, the response-aligned, trial-averaged CPP waveforms exhibit modulations as a function of evidence strength or RT that are widely reported in previous research and interpreted as empirical support for the CPP’s role in sensory EA. However, when a deconvolution method was applied, which models the ERP as the sum of stimulus-locked (S) and response-locked (R) components that overlap according to RT, pre-response amplitude effects seen in the original waveforms no longer reached statistical significance after the S component had been removed from the data. From this the authors conclude that the CPP’s pre-response amplitude modulations were an artifact of averaging the overlapping S and R components across trials. They go on to suggest that this observation calls into question the CPP’s purported role in tracing sensory EA.

Here, we argue that the analysis and interpretation of these perceptual choice data are critically flawed. We first demonstrate with simulations that the deconvolution analyses used by the authors are not designed to correctly capture bounded accumulation signals. Applying these analyses to true, ramping, accumulation-to-bound signals can result in the same amplitude effect reductions as in Frömer et al’s empirical results. Second, we highlight key empirical results that are misinterpreted by the authors. Finally, while Fromer et al focus on pre-response amplitude modulations by RT and difficulty, we highlight the diversity of other EA signatures that have been identified in the CPP in the extant literature which cannot be explained by simple signal overlap.

### Unfold mischaracterises evidence accumulation signals

The Unfold toolbox ^3^ implements deconvolution (i.e. overlap-correction) under the assumption that the only signals present in the EEG are neural responses precisely time-locked to either the stimulus or response (Fig 1a). Indeed, in this situation the RT-dependent summation of the S and R components can cause false RT effects in trial-averaged response-locked data even when there is no RT effect in the underlying ground-truth components. This ground-truth is recovered after applying Unfold (Fig 1c; recapitulating Frömer et al Figure 5). However, the accumulation-to-bound processes invoked in sequential sampling models are fundamentally different. In standard accumulation models that assume stationary evidence, the accumulation process begins at (or shortly after) the stimulus and ends at response. In this way it is neither stimulus-locked nor response-locked, but rather interposed between the two, temporally stretching/contracting in slow-/fast-RT trials (Fig 1b). There is currently no means to capture such timescale-variation directly in the Unfold toolbox. To demonstrate the impact of this, we applied the same Unfold analyses as were applied to the contrast discrimination dataset ^4^, to a simulated S-R-interposed ramp signal, which minimally represents the core element of accumulation-to-bound processes ^5–7^. The Steinemann et al task requires an immediately-reported decision about a single, stationary sensory feature (contrast difference), whose accumulation would approximate a straightforward, S-R-interposed ramp on average.

**Figure 1:**
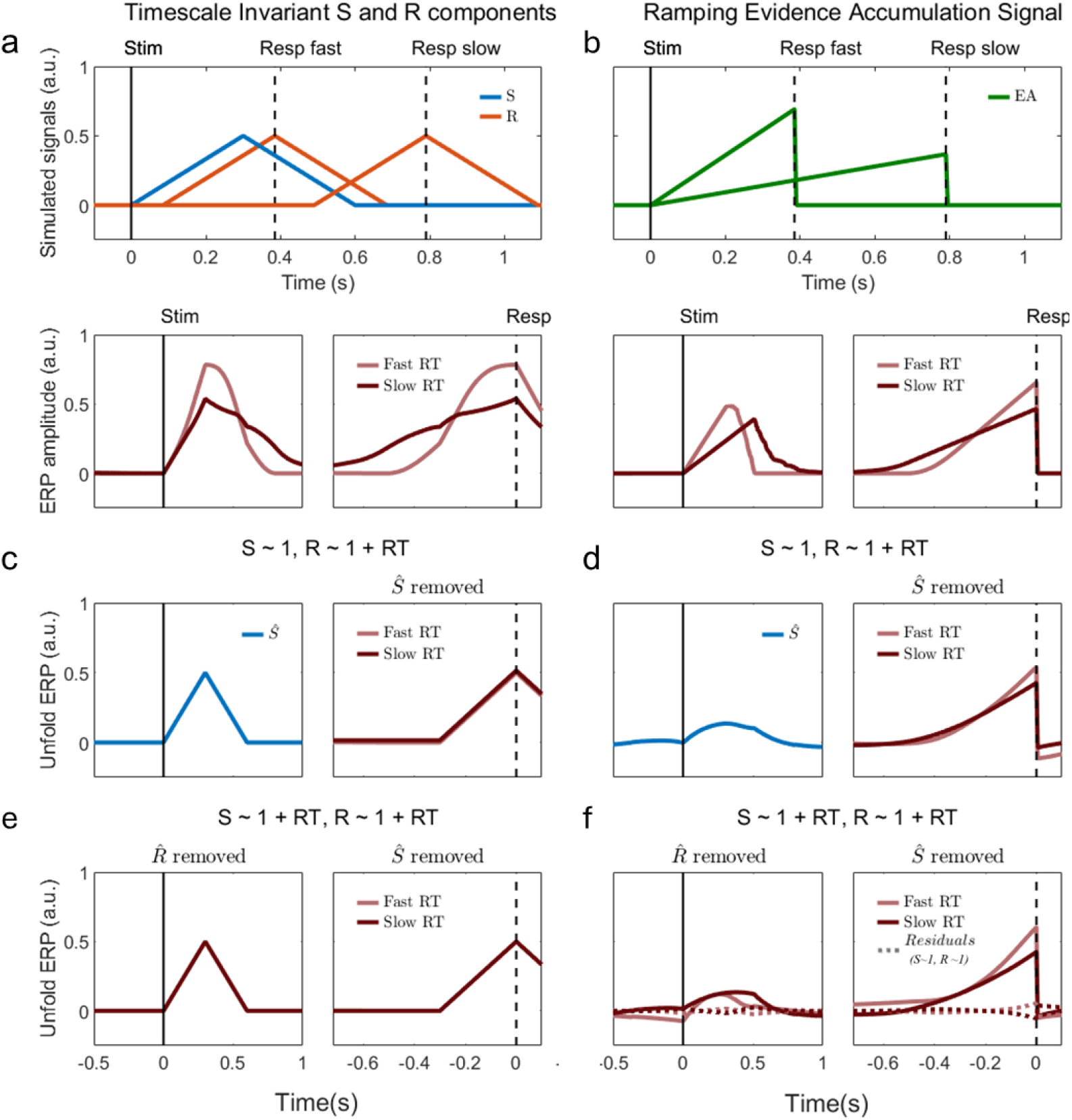
Outcomes of using Unfold on simulated data that do (timescale-invariant S and R components), versus do not (S-R interposed ramp), adhere to the underlying assumptions of Unfold. **A)** We generated ground-truth simulated data with only timescale-invariant S and R components, and no effect of RT on the amplitudes of either the S or R component. Two example trials from this scenario are shown, one around the 10th (Resp fast) and one around the 90th (Resp slow) RT percentile (top). Trial-averaged timecourses aligned to stimulus and response show that RT-dependent overlap causes RT effects in the response-locked traces (bottom). **B)** In an alternative simulated scenario, there are no S or R components but rather a single accumulation-to-bound process ramping from stimulus to response, with a collapsing bound (top). Trial-averaged traces (bottom) show that in this ground truth, there are effects of RT on both the buildup rate and amplitude reached at response when average traces are split by median RT (e.g. as observed in^4^ &^8^. **C)** In the timescale-invariant S & R simulations, “correcting” the signal by removing the unfold-estimated S-component (blue) from each trial leaves a response-aligned waveform that precisely matches the R-component created, with no RT effects. **D)** However, running the same analysis on a ramping evidence accumulation signal spuriously allocates some of the initial part of the ramp to the S component (blue) whose subtraction dramatically reduces the real RT effects on both slope and amplitude in the response-aligned trace, as observed in Frömer et al (see their Figure 6). This spurious reduction in RT-dependent buildup occurs for constant bounds also; we simulated a collapsing bound here to produce qualitatively similar patterns as in the ‘uncorrected’ contrast discrimination data of Steinemann et al^4^. Next, Frömer et al sought to test whether, even if reduced from the response-locked waveforms, the RT effects may appear instead in the stimulus-locked waveforms after removing the R components. Whereas in their main analysis only the R component was allowed to vary between fast and slow RT categories, here they allowed RT effects in both the S and R components. They found that stimulus-locked waveforms for fast- and slow-RT trials had similar peak amplitudes, but with a large RT effect (fast < slow) now appearing in the baseline period (their Supplementary Figure 6C). **E)** When this analysis was applied to the time-invariant S-R component simulations, no such baseline shift was found, and Unfold perfectly recovered the original signals. **F)** In contrast, the simulated ramping evidence accumulation signal exhibited the same baseline shifts observed in the authors’ analysis of the Steinemann et al data. Note that the degree and morphology of such artifacts arising from Unfold depend strongly on analysis settings (e.g. time-expansion window, regression structure, etc), and while we have attempted to match those applied to the Steinemann et al dataset here, the code that was shared with the paper indicates that other perceptual decision datasets were analysed with different settings. Finally, Frömer et al reported that removing both the R and S components of an RT-agnostic regression leaves very little residual activity in the empirical ERP data (see their Supplementary Figure 7), which they interpret as evidence that Unfold has not removed real EA activity. Again however, we observe qualitatively the same outcome when this subtraction is performed on the simulated ramp data (dashed traces in panel F). Thus, systematic misinterpretations arise from the fact that these Unfold analyses do not directly test for the presence or absence of an accumulation signal. All of the same outcomes are observed when these analyses are conducted on a simulated diffusion-type EA process (see Supplementary Figure 1).

As expected, the Unfold analysis spuriously ascribes an initial portion of the S-R-interposed ramp to the S component and the end portion to the R component (Fig 1d). In so doing, Unfold systematically blunts RT effects that are truly present in the data, because the inappropriate subtraction of a spurious S component reduces the amplitude of the fast-RT waveform close to response while reducing the slow-RT waveform further back in time. Thus, while Unfold does correctly eliminate RT effects where they falsely arise from S- and R-component overlap, it also erroneously removes RT effects when they are truly present in a ramping accumulator signal.

### Signatures of evidence accumulation remain evident after deconvolution with Unfold

Fromer et al themselves provide simulations of an EA process with additional features such as diffusion noise and non-decision time delays (their Supplementary Figure 5). When discussing these, Fromer et al focus on the fact that, while Unfold does divide the EA activity between the S and R components, none of the EA activity is lost, as one could accurately reconstitute the original signal from the estimated S and R components. However, this appraisal overlooks the key fact - clearly visible in their simulations - that when spread across separate S and R component estimates, true RT effects are greatly weakened so that they are bound not to be detected by statistical tests applied to one component at a time.

Frömer et al also use the simulations to show that the more latency jitter in the post-accumulation (motor) delay relative to pre-accumulation (stimulus encoding/transmission) delay, the more a true EA signal is attributed to the S component, and even in this case, its amplitude is larger for fast compared to slow RTs (right panel). From this, the authors conclude that, if Unfold has in fact attributed EA activity to the S component of the real EEG data, then it should exhibit a similar amplitude modulation. When the authors tested this, of the two datasets that exhibited the relevant amplitude modulation in their uncorrected stimulus-locked waveforms (visual search and contrast discrimination), both also showed those modulations in their S components after deconvolution (Supplementary Figure 6b&c). In the case of the contrast discrimination data, deconvolution causes the RT effect on amplitude to transfer to the pre-stimulus baseline period, so that the peak amplitude effect would emerge with baseline-correction, as noted by Frömer et al. As our Figure 1f shows, the same spurious pre-stimulus amplitude difference can arise as an artifact of applying Unfold to an EA signal. Based on the observation that little residual activity remains after subtracting both the S and R component estimates from the empirical data, Frömer et al conclude that no EA signatures are present in the contrast discrimination data (Supplementary Figure 7). However, our simulations show that the same outcome is observed for a true underlying EA signal, whether in the form of a simple ramp (Figure 1f), or a more elaborated diffusion process (Supplementary Fig1).

In general, the simulations in their Supplementary Figure 5 show that the degree to which any EA signal will be aligned to the final response depends on the relative amount of pre-versus post-EA latency jitter. This, in turn, is likely to be highly task-dependent. For example, pre-EA variability can be expected to increase in tasks in which the location of the physical evidence is not known in advance while post-EA variability will increase with the complexity and speed of the decision-reporting actions. However, while these factors can change the temporal duration of EA as a proportion of the total RT, none of them change the very fact of it being an evidence accumulation process.

### Various signatures of EA have been identified in the CPP without recourse to analyses of response-aligned average waveforms

Whereas Frömer et al focus on deconvolution analyses to interrogate amplitude modulations in trial-averaged waveforms, their discussion of the previous literature overlooks the many alternative approaches to isolating signatures of EA employed in previous studies of the CPP. These include designing stimuli to eliminate sensory-evoked potentials at evidence onset, detailed analysis of single-trial signal time courses, application of current source density transformations to reduce spatial spread due to volume conduction, hypothesis-driven experimental manipulations and model-based analyses (reviewed in ^2^).

This work has identified numerous CPP features that are consistent with EA and which cannot be readily attributed to component overlap. Examples include, but are not limited to: single-trial surface plots that exhibit a progressive build up extending from shortly after evidence onset until the perceptual choice report ^9^; ^10^; ^11^; after statistically controlling for RT, single-trial measurements of the CPP’s pre-response amplitude are modulated by experimental manipulations of time pressure in a manner that accords with the decision bound adjustments identified by sequential sampling models ^4,8^; the pre-response amplitude of the CPP is larger on trials in which the correct alternative was invalidly cued despite slower RTs also in accordance with model-identified boundary adjustments ^8^; ^12^; subtle perturbations ^11^ or random fluctuations ^13^ of the physical evidence while decision formation is ongoing leads to corresponding variations in the CPP’s build-up in stimulus-aligned measurements that predict subsequent behavioral effects; both stimulus- and response-aligned single-trial measurements of the CPP’s build-up rate are correlated with choice confidence after controlling for RT, accuracy and difficulty ^14^; the CPP’s stimulus and response-aligned average timecourse, as well as modulations of its pre-response amplitude as a function of RT, evidence strength, accuracy, time pressure and prior knowledge, can all be recapitulated by simulating cumulative evidence signals from a sequential sampling model that provides excellent behavioral fits ^8^.

If considered only one at a time, some of the above results could potentially be explained by invoking mechanisms other than sensory EA. However, sensory EA is the only account we are currently aware of that can accommodate all of the above observations at once and in detail. Finally we note that the CPP features observed even in the value-based choice data of Frömer et al are also consistent with it tracing the perceptual decisions (i.e. stimulus identification) that are a prerequisite for initiating value comparisons in this task.

These features include the fact that its peak falls well before the final response (the value choice process can only start once the stimuli have been identified), that its peak falls closer to the final response on trials with faster RTs (faster value choices will cause the response-locked ERP to overlap more with the perceptual choice process), that Unfold assigns it to the S component and that, once isolated in the S component, it no longer exhibits any value choice-related modulations. While the above observations accord with the CPP tracing perceptual decisions for the value task, an interesting challenge for future research will be to determine whether those decisions take the form of an EA process.

In summary, we show that the regression-based deconvolution method used in Frömer et al. (2024) is unsuitable for correctly identifying and characterizing EA signals. There is also extensive prior evidence that the CPP traces sensory EA, and this evidence is not dependent on analyses of response-locked ERPs. Thus, the authors’ claims relating to the CPP and sensory EA are not supported by their data. Component overlap is undoubtedly an important issue to account for in noninvasive neurophysiology, and to the extent that paradigm design cannot fully eradicate them ^15^, analysis tools such as deconvolution can in principle be helpful; however, as the above discussions highlight, to be used for this purpose the deconvolution methods must be endowed with the ability to capture EA signals and characterise them correctly as EA signals, and at minimum this should involve components whose timescale can vary with RT ^16^.

## Acknowledgments

The authors would like to thank Romy Frömer, Benedikt Ehinger, Matthew Nassar and Amitai Shenhav for their feedback on this commentary and for many useful discussions relating to their paper. We also thank our laboratory teams as well as Michael Nunez, Kobe Desender, Martin Wiener, Laurence Hunt, Lucas Parra and Jaeger Wongtrakun for helpful comments. This work was funded by an SFI research grant to R.G.O. and S.P.K. (19/US/3599). R.G.O. was funded also by the European Research Council Consolidator Grant IndDecision – 865474, and S.P.K. also by a Wellcome Trust Investigator Award (219572/Z/19/Z). D.F. was supported by an Australian Research Council Discovery Early Career Researcher Award (ARC DE220101508). E.A.C. was funded by the European Research Council Starting Grant Myodecision (101077772).

---

**Supplementary Figure 1.**
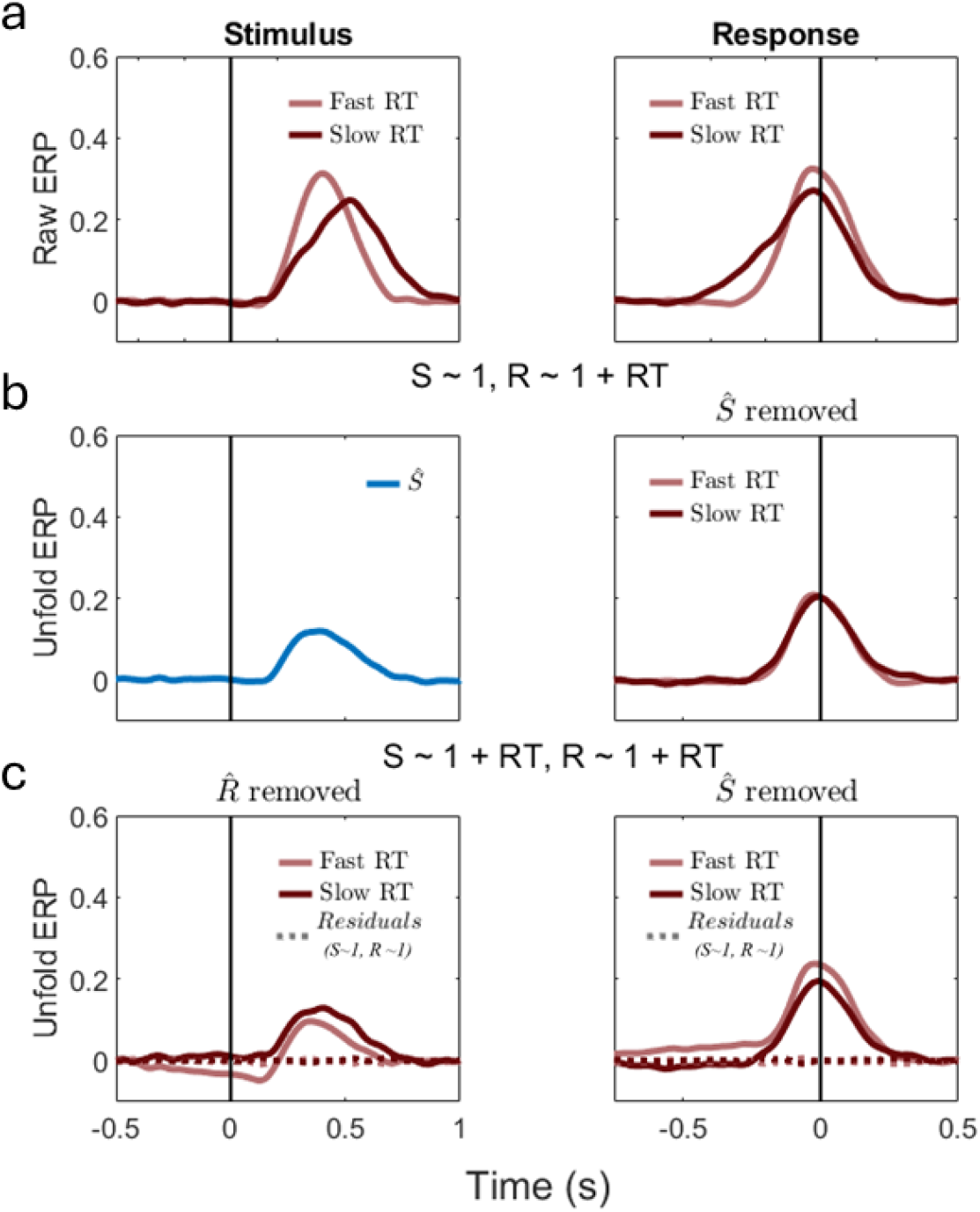
Replication of all analyses reported in Figure 1 but this time applied to a noisy diffusion process simulated from the same code used to generate Frömer et al’s Supplementary Figure 5. **A)** Trial-averaged timecourses aligned to stimulus and response exhibit RT-dependent effects on both the buildup rate and amplitude reached at response. **B)** Unfold assigns a substantial portion of the EA signal to the S component (left panel). Removal of that S-component dramatically reduces the real RT effects on both slope and amplitude in the response-aligned trace, as observed in the empirical data analyses of Frömer et al (see their Figure 6). **C)** Spurious reduction of effects was also observed when allowing both the S and R components to vary with RT. Removing the estimated S and R components from an RT-agnostic regression produces virtually flat residuals, as observed in the empirical data analyses of Fromer et al. (their Supplementary Figure 7). Whether or not residual activity consistent with an evidence accumulation process can be found after applying unfold may be dependent on specific data features (e.g. RT distribution, stimulus/motor variability, etc) and Unfold settings. However the key message of this simulation is that Unfold can result in flat residuals even for a ground truth EA signal, and therefore Frömer et al’s results are not informative regarding the presence or absence of EA signals in their empirical data.

